# BlastFrost: Fast querying of 100,000s of bacterial genomes in Bifrost graphs

**DOI:** 10.1101/2020.01.21.914168

**Authors:** Nina Luhmann, Guillaume Holley, Mark Achtman

## Abstract

BlastFrost is a highly efficient method for querying 100,000s of genome assemblies. It builds on Bifrost, a recently developed dynamic data structure for compacted and colored de Bruijn graphs from bacterial genomes. BlastFrost queries a Bifrost data structure for sequences of interest, and extracts local subgraphs, thereby enabling the efficient identification of the presence or absence of individual genes or single nucleotide sequence variants. Here we describe the algorithms and implementation of BlastFrost. We also present two exemplar practical applications. In the first, we determined the presence of the individual genes within the SPI-2 *Salmonella* pathogenicity island within a collection of 926 representative genomes in minutes. In the second application, we determined the existence of known single nucleotide polymorphisms associated with fluoroquinolone resistance in the genes *gyrA, gyrB* and *parE* among 190, 209 Salmonella genomes. BlastFrost is available for download at https://github.com/nluhmann/BlastFrost.

## 1 Introduction

Recent advances in DNA sequencing technologies have reduced sequencing costs and time, and whole-genome sequencing of bacterial pathogens is being routinely perfomed by public health organizations. The resulting sequence reads and genome assemblies are deposited in the public domain [3, 25, 30], enabling comparative analysis for 100,000s of genomes [17, 31] from individual bacterial genera to investigate their evolutionary history, and their assignment to ongoing or historical epidemiological outbreaks.

New sequencing data are now routinely uploaded to public databases such as the Sequence Read Archive (SRA [16]), which has resulted in ready access to extensive collections of sequencing data for many bacterial genera. Other databases, including PubMLST [15] and EnteroBase [30], even provide curated collections of genomic assemblies bundled with their metadata for specific bacterial pathogens, as well as dedicated tools for population genomic analyses.

The analysis of genomic sequences by phylogenetic approaches can yield insights into evolutionary distances for 1000s of genomes but large comparative studies based on sequencing data are limited by computing resources and speed of calculations [31]. Even the seemingly simple task of identifying all bacterial strains within a collection that contain a specific antimicrobial resistance gene or other genes of interest is a computational challenge for the large data sets that are currently available. The most popular methods for sequence comparison are BLAST [5] and its successors. However, these alignment-based methods do not scale well. As a result, in some recent software the alignment step is replaced by a *k*-mer approach, in which sets of short sub-sequences of fixed length *k* are compared between a query and a sequence database, as recently reviewed by Marchet et al. [20]. Without the prerequisite for an explicit reference genome, *k*-mers can identify diverse genetic modifications such as SNPs (single nucleotide polymorphisms), insertions or deletions from short read sequences.

One recent *k*-mer-based method, BIGSI, employs a data structure storing a Bloom filter [6] of k-mers for each genome in a database, and can subsequently index and search very large databases of bacterial and viral sequences [7]. BIGSI queries are very efficient, but the 2016 European Nucleotide Archive (ENA) is so large that creating a BIGSI index took months. Furthermore, BIGSI was designed for dealing with genetically diverse collections of data, and other methods and different data structures might be more efficient for creating a query index of sequence data from closely related genomes. One potential approach for faster construction of indices would be to index sets of *k*-mers in a de Bruijn graph [24], where shared k-mers are automatically collapsed. Collapsing *k*-mers that are shared between closely related genomes would decrease both the storage space for the index and the search space for subsequent queries. Recent implementations of such an approach include Mantis [23], Rainbowfish [4] and VARI-Merge [22]. They build joint de Bruijn graphs for multiple genomes, coloring nodes by their source genomes (colored de Bruijn graphs [13]), and can traverse shared paths in the graph which represent conserved regions as well as diverging paths which represent variable regions. However, the implementations of these methods do not scale well enough to be able to handle a modern, large sequence collection [12]. For example, VARI-Merge was benchmarked on a data set of 16,000 Salmonella genomes [22], but EnteroBase already contains ∼ 250, 000 genomes.

The recent development of Bifrost [12] introduced a memory efficient, dynamic data structure for indexing either colored or non-colored compacted de Bruijn graphs. It presents a broad range of functions that support querying both sequences and colors, annotating individual vertices and editing Bifrost graphs while preserving their compaction. The implementation of Bifrost facilitates to rapidly build joint graphs scaling to 100,000s of genomes, and permits almost instantaneous updating of graphs with additional data. However, Bifrost does not implement querying on its own. Here we introduce BlastFrost, a method for similarity searches in Bifrost graphs by rapid *k*-mer matching implemented in C++. BlastFrost can use the underlying Bifrost graph structure to extract subgraphs defined by a query, and thereby efficiently extract sequence variants of the query from a data base of 100,000s of bacterial genomes, as illustrated here by case studies on the identification of genomic islands and of individual mutations in antimicrobial resistance genes.

## 2 Results

Bifrost indexes bacterial genomes in a time and memory efficient implementation of a compacted and colored de Bruijn graph. The nodes in an uncompacted graph represent a set of overlapping sequences of k-mers within the input genomes. Edges in the graph are implicit, and represent overlaps of length *k* − 1 between neighbouring nodes. Maximal paths of multiple sequential, non-branching nodes are compacted into single nodes (unitigs) by collapsing the overlaps. Each node is assigned a set of colors representing all input genomes containing the corresponding k-mers of the unitig.

BlastFrost relies on the particular form of compacted and colored de Bruijn graphs implemented in Bifrost [12], which are henceforth designated as Bifrost graphs. As depicted in Figure 1 top, we implemented a k-mer search function in BlastFrost which can identify the presence or absence of a query sequence in any of the genomes in a Bifrost graph. The results of that search can be used for subgraph extraction (Fig. 1 bottom) of query matches in order to identify all variants of the query sequence in the Bifrost graph. The following paragraphs provide an overview of the method. Algorithmic details can be found in Supplemental Material.

**Figure 1:**
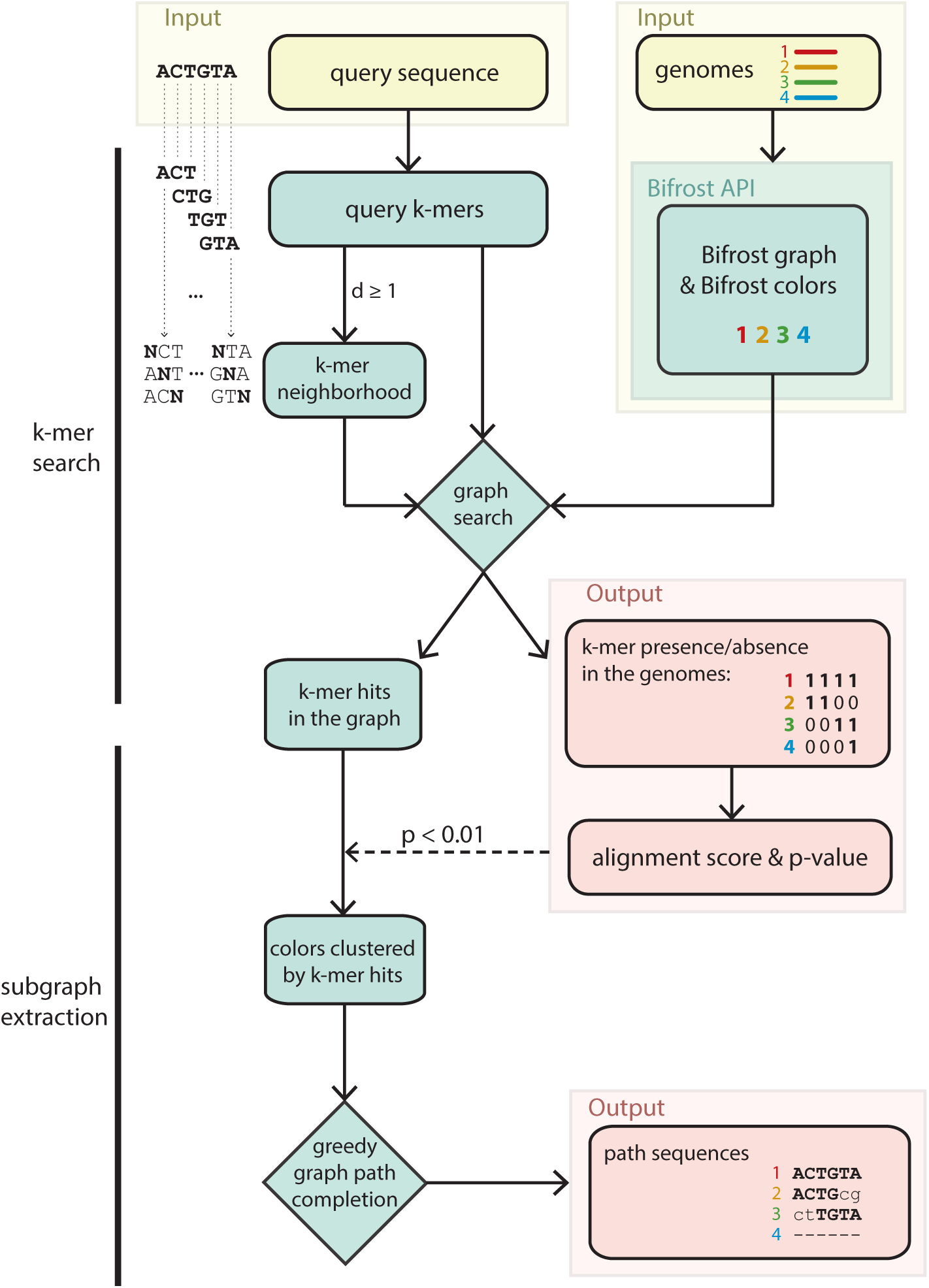
BlastFrost algorithms. BlastFrost is a command line program with inputs of pre-computed Bifrost files that specify the graph and colors plus a FASTA file of query sequences. It conducts a search of a *k*-mer neighbourhood for the parameters *k* (*k*-mer length) and *d* (Hamming distance), and estimates an alignment score and p-value for each query sequence. The presence or absence of hits are automatically produced as a tab-delimited file containing the query ID, color ID, and binary presence/absence data. When started with the input parameter −*e*, BlastFrost uses these data to extract subgraphs, and adds their path sequences as output.

### 2.1 BlastFrost query search

BlastFrost takes as input, and loads into memory, a pre-computed Bifrost graph for a certain *k* consisting of a graph file in GFA format plus an index of the colors of each *k*-mer in each unitig. We henceforth refer to the graph genomes as colors. The input parameters to BlastFrost also include a link to a FASTA file containing one or more query sequences.

For each query, BlastFrost calculates a set of overlapping k-mers whose size corresponds to the value of k that was used to build the Bifrost graph. That set is used to identify the presence or absence of sequences corresponding to that query in all colors. BlastFrost searches for each *k*-mer via functions that are integrated in the Bifrost API. Each query results in a binary sequence for each color of 1s and 0s representing k-mer hits and misses. Note that the *k*-mer based search in the graph explicitly assumes that any two overlapping k-mers of the same color are also contiguous in the underlying genome. Since Bifrost graphs are compacted, BlastFrost additionally speeds up comparisons for the compacted nodes in the graph (unitigs), by assuming that the color set of the unitig is the same as the individual color sets of each k-mer in that unitig.

A single nucleotide substitution between a query and a color will result in *k k*-mers that are missed for that query, resulting in a stretch of k 0’s in the binary hit sequence for that query. Deletions are also characterized by runs of 0’s that are potentially smaller than *k*, while insertions and multiple substitutions can lead to longer runs of 0’s in the hit sequence. In order to evaluate the significance of *k*-mer hits between a query and a specific color, we adopt the BLAST approach for computing an E-value based on an estimated alignment score, derived from the lengths of 0 runs in the *k*-mer hits. To increase the sensitivity of the *k*-mer based query, BlastFrost allows additional querying of all *k*-mers related to a query *k*-mer by a Hamming distance smaller or equal to an input parameter *d*. We refer to this set of additional *k*-mers as *k*-mer neighbourhoods (Fig. 1). In the following evaluation, we present the necessity for this increased sensitivity, as well as some of the resulting trade-offs.

### 2.2 BlastFrost subgraph extraction

Presence or absence results from a Bifrost graph are not immediately informative on genomic location of the query hits, the numbers of copies in each genome, or on syntenic relationships. We note that for any specific query, each binary sequence of *k*-mer hits represents a potentially incomplete path of nodes for each color in the graph interrupted by nucleotide changes that were not included in the shared *k*-mers between the query and the genome. BlastFrost can account for these potential gaps by extending the k-mer hit results, and produce a subgraph for each successful k-mer query. Starting from the first unitig in the original k-mer hit list for a specific color, BlastFrost greedily completes a path by traversing non-branching paths of the same color within the graph. The subgraph is then used to reconstruct the corresponding sub-sequence of each color from the path in the Bifrost graph.

To avoid completing the same paths multiple times, BlastFrost clusters colors sharing k-mer hits in order to simultaneously complete all their paths, and also removes colors from those clusters that are absent in intervening unitigs. For each path and its accompanying colors, BlastFrost output the genome sequence in addition to the above mentioned binary sequence. These data allow ready identification of variant positions between the query and the extracted path sequences.

### 2.3 Evaluation and benchmarks

#### Precision and sensitivity of identifying all query variants in a pangenome

We evaluated the abilities of BlastFrost for detecting sequence variants within a Bifrost graph by querying all 21, 065 orthologs in the whole genome MLST (wgMLST) scheme in EnteroBase [2, 30] which were derived from a pan-genome from 537 representative genomes of the genus *Salmonella*. Bifrost required less than 24 minutes and less than 5GB of memory to create a graph of 926 representative *Salmonella* strains from EnteroBase [2]. The graph requires 2.3GB of disk space, and it contains more than 33 million unitigs. We queried the Bifrost graph with BlastFrost for one representative allele from EnteroBase for each of the 21, 065 loci in the wgMLST scheme, and extracted all allelic variants in the corresponding subgraphs. For each locus, a allelic variant was scored as successfully recovered if at least 95% of the allelic sequence stored in EnteroBase was present within the BlastFrost output.

Figure 2 depicts the sensitivity and precision of this evaluation by sequence length for pairs of queried and recovered alleles according to their average sequence identity. BlastFrost identified all genomic alleles which were at least 95% identical on average to the query (100% sensitivity). For more diverse alleles, the sensitivity dropped to > 0.89 for sequences that were less than 400bp long. The number of false hits was very low (precision > 0.9) for alleles longer than 400bp, but somewhat more common for shorter sequences. It is not clear that these false hits are really false, because EnteroBase filters repetitive DNA and overlapping or duplicated alleles from its allelic calls whereas those sequences are still present in the genomes, and can be found by BlastFrost. In summary, BlastFrost correctly identified all sequence variants down to 90% sequence identity with a query length of > 400bp, and almost all such variants down to 200bp in length.

**Figure 2:**
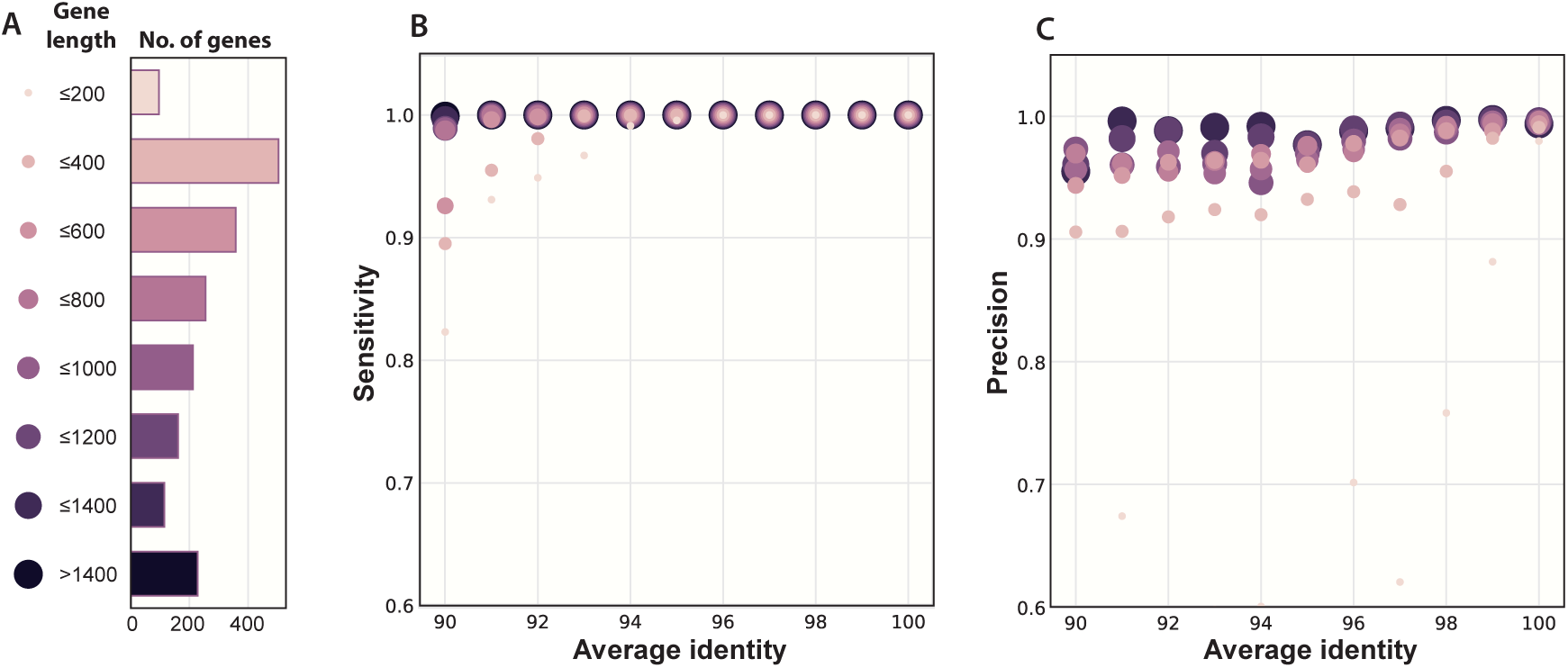
Sensitivity and precision of extracted subgraph sequences of 21, 065 wgMLST loci in the Salmonella database in EnteroBase using their reference alleles as queries against a Bifrost graph of 926 representative *Salmonella* strains [2]. Queries were scored as correct when at least 95% of the sequence length of the reference was retrieved and EnteroBase had scored the genome as containing an allele of that locus. Query hits that were retrieved from genomes which were scored as missing that locus by EnteroBase were scored as false positives. In contrast, loci reported as present in that genome by EnteroBase which did not result in a BlastFrost hit were scored as false negatives. A). Distribution of numbers of wgMLST loci by sequence length. B) Sensitivity. C) Precision.

#### Speed and RAM benchmarking

Time and memory requirements were compared against two widely used, recently developed software tools. Firstly, BlastFrost was compared to BIGSI [7] on a data set of 736 representative *Yersinia pestis* draft genomes, all of which are very closely related genetically. Bifrost indexing was much faster than that of BIGSI (Supplemental Material). Figure 3 shows query time and maximum RAM usage of BlastFrost and BIGSI as a function of the size (200-1600) of subsets of core genes from the EnteroBase core genome MLST (cgMLST) scheme for *Yersinia* [30]. The BlastFrost search was an inexact search for a *k*-mer neighbourhood of 1 (parameter *d* = 1). The exact search function in BIGSI was tested as well as its inexact search function which reports query hits containing at least 70% of the query sequence k-mers (parameter *t* = 70). The BlastFrost query yielded the same hits as the inexact BIGSI search, but was as fast as the exact BIGSI search (Fig. 3A), and used much less RAM except when > 1200 queries were performed.

**Figure 3:**
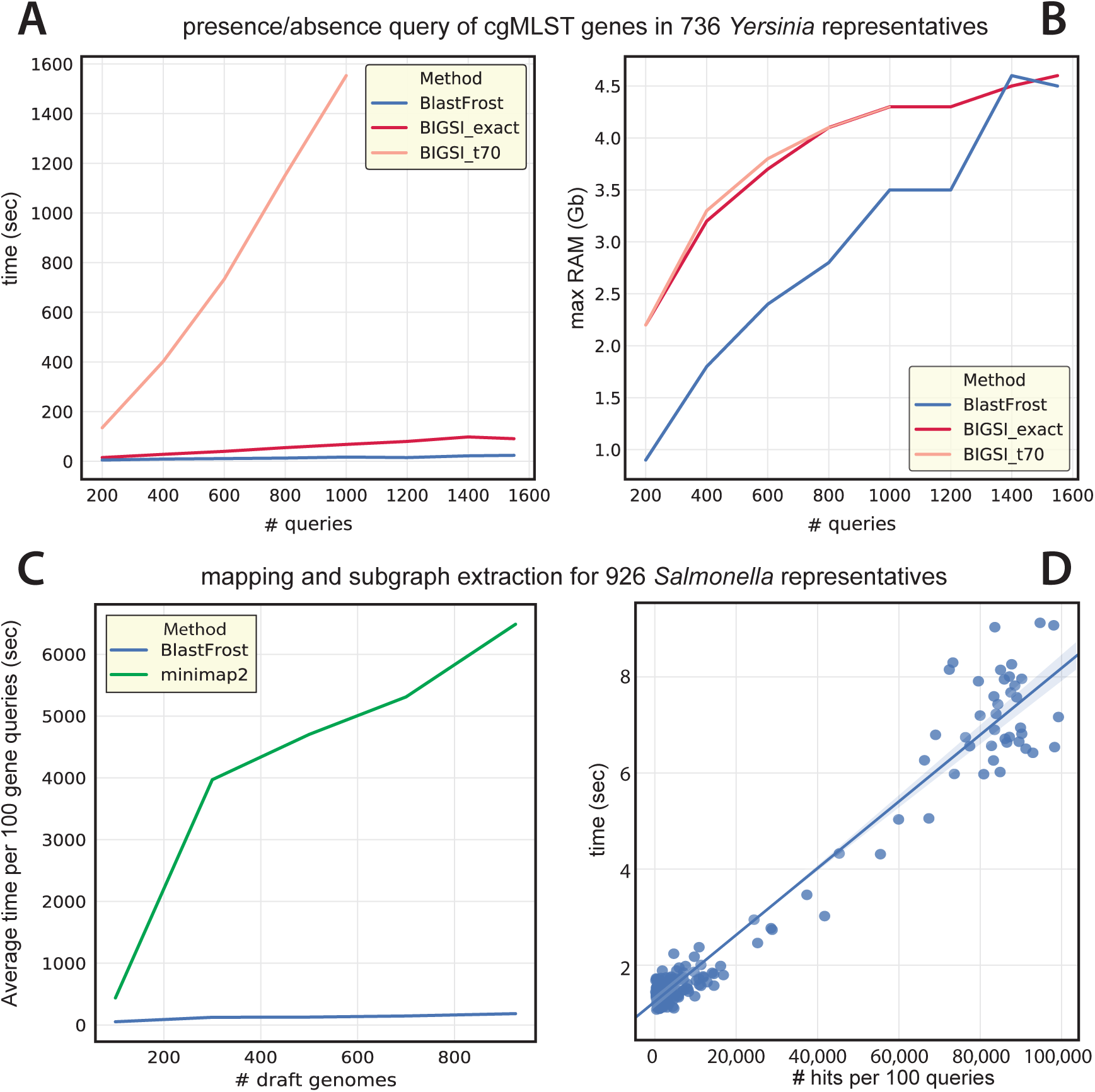
Speed and memory of BlastFrost for cgMLST genes in comparison to BIGSI (A,B) and minmap2 (C) and as a function of numbers of hits (D). (A, B) Speed and memory with 736 genomes of *Yersinia pestis* [30] as a function of numbers of queries for BlastFrost with *k*-mer neighbourhood *d* = 1 in comparison to both exact and inexact BIGSI queries. C) Speed with subsets of 926 *Salmonella* genomes [2] as a function of numbers of draft genomes for BlastFrost with *k*-mer neighbourhood *d* = 1 in comparison to minimap2. (D) Correlation between time required by BlastFrost to extract a subgraph by number of hits for a 100 gene query.

For the speed and RAM needed for subgraph extraction, we compared BlastFrost to minimap2 [18], which is the currently most efficient mapping tool for both short reads as well as chromosome-scale alignments. The average speed of the two methods was tested for extracting 100 genes at a time from the wgMLST *Salmonella* scheme described above from subsets of the 926 representative *Salmonella* genomes. Figure 3C shows much higher speed for BlastFrost than minimap2. The time needed by BlastFrost to extract a subgraph is dependent on the number of hits for that query (Fig. 3D), but it still achieves a slightly sub-linear growth in time requirement because several genomes can share a common sequence for the same query variant within a bacterial genus.

### 2.4 Applications

We took advantage of the large genomics databases available in EnteroBase to demonstrate the ability of BlastFrost to find the presence of genomic elements and to identify nucleotide variants of individual genes. For genomic elements, we searched the representative genomes of Salmonella for known genes in the SPI-2 *Salmonella* pathogenicity island. For nucleotide variants, we screened the entire EnteroBase *Salmonella* database for specific substitutions in three genes that are associated with fluoroquinolone resistance in *Salmonella*.

#### Genomic islands

Genomic islands consist of DNA stretches in the accessory genome that can be acquired by bacteria through horizontal gene transfer, or which are lost due to gene deletion [8, 27]. Pathogenicity islands are a distinct class of genomic islands, which can range in size from 10 − 200kb, and encode genes which can contribute to the virulence of the bacteria [11]. SPI-2 is such an island which seems to have been acquired by *Salmonella* after the divergence of *S. bongori* and *S. enterica* from their common ancestor. Subsequently *S. enterica* split into multiple so-called sub-species [2].

We obtained gene sequences from the Virulence Factors database [19] for the 44 genes on SPI-2in *S. enterica* serovar Typhimurium strain LT2, and queried each of these against the same Bifrost graph of the 926 representative *Salmonella* genomes described above. Figure 4 shows the distribution of these 44 genes according to an exact search (BlastFrost parameter *d* = 0, dark green) and an inexact search (BlastFrost parameter *d* = 2, light green). The inexact search consistently identified most of the SPI-2 genes in all of the *Salmonella* subspecies, but they were absent, as expected [11], in *Salmonella bongori*. However, some genes were absent or their sequences were too divergent to be detected from most of the genomes from individual subspecies. This figure also emphasises the importance of inexact querying because although most SPI-2 genes in subspecies I, II and VI can be identified by an exact search, the inexact search greatly increased the number of SPI-2 genes identifiedin the other subspecies.

**Figure 4:**
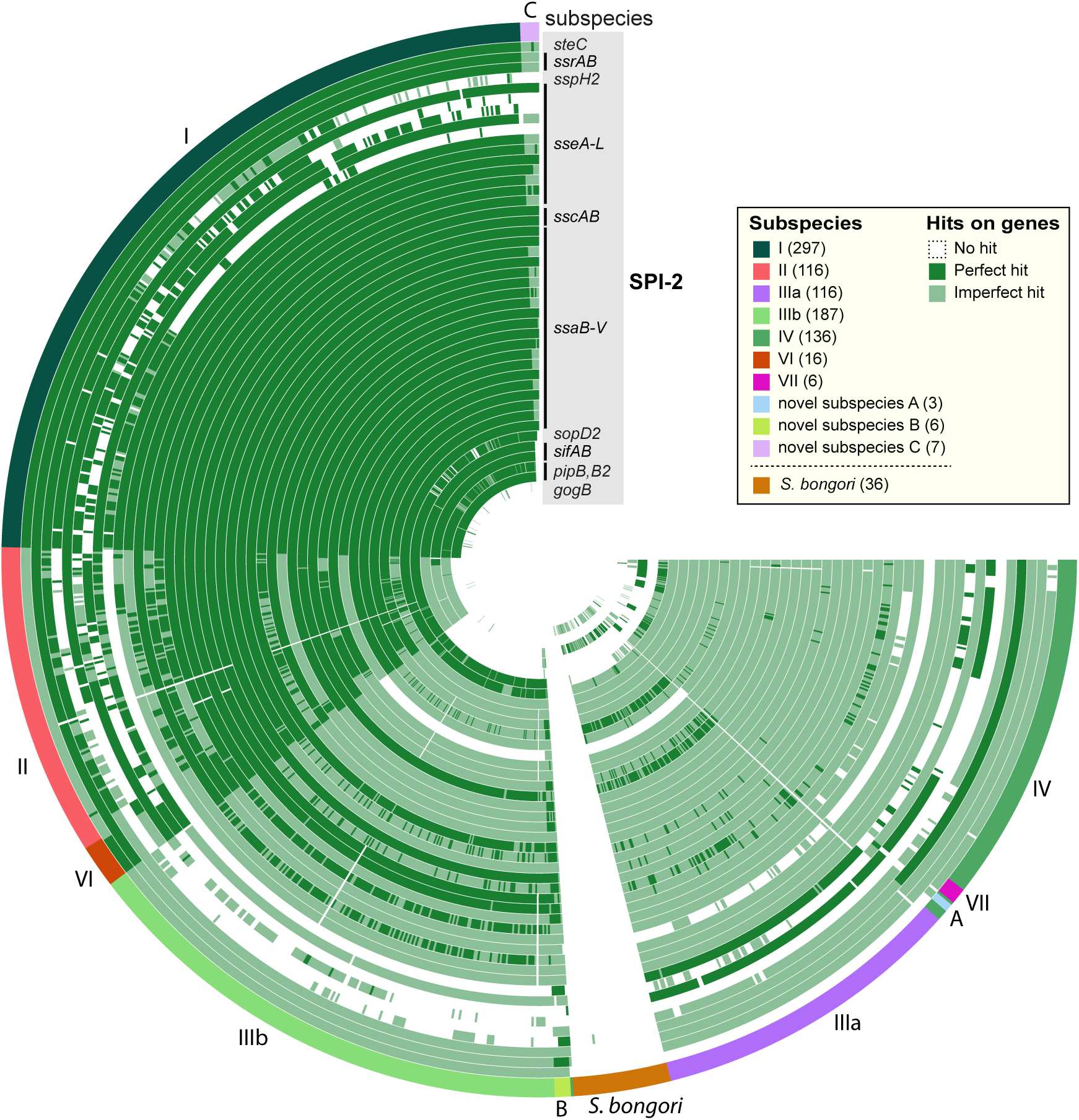
BlastFrost presence/absence analysis for 44 genes of the Salmonella SPI-2 pathogenicity island among 926 representative *Salmonella genomes* [2]. Dark green indicates hits identified by an exact (*d* = 0) or inexact (*d* = 2) search while light green indicates hits identified only by the inexact search. White indicates no hits. Each concentric circle shows the results for the gene indicated in the figure legend at the top right. The circle consists of arcs, with one segment for each genome from *Salmonella bongori* and the subspecies of *Salmonella enterica* that are indicated outside the circles. Graphical representation was with Anvi’o [9].

This analysis took 111 seconds to load the Bifrost graph of 926 *Salmonella* genomes into memory, and a further 540 seconds to search for all SPI-2 genes with the inexact BlastFrost search, for a total of under 11 minutes.

#### Nucleotide variants

The subgraph extraction functionality of BlastFrost can also extract known variants of genes involved in antimicrobial resistance or other phenotypes. We initially created a Bifrost graph of 160, 000 *Salmonella* draft assemblies downloaded from EnteroBase, which took 4 days and 15*h* computation time and 147GB of memory. During the course of these investigations, we updated this graph in several iterations, resulting in a final graph containing 190, 209 genomes. A Bifrost graph update with 100 additional genomes takes about 2.5h, including the time to load the graph back into memory. The disk size of the graph containing 190, 209 genomes is 158.5GB and it contains 32, 692, 889 unitigs. We then queried this graph for a single representative gene sequence from each of the genes *gyrA, gyrB* and *parE*. These genes were chose because individual nucleotide variants in the quinolone resistance-determining regions (QRDR) can cause reduced susceptibility to fluoroquinolones [26]. The queries resulted in one subgraph per gene, whose sequences were aligned, and scanned for the known nucleotide variants.

Our results show that 20, 490 genomes from multiple serovars contained these QRDR nucleotide variants, as illustrated by a Neighbour-Joining tree estimated from cgMLST distances between the genomes colored by serovar (Fig. 5A). The serovars of genomes containing QRDR mutations included common causes of human disease, such as Enteritidis, Typhi and Typhimurium, as well as multiple other serovars that are common in domesticated animals but can cause food-borne gastroenteritis in humans (Fig. 5B). Most of the genomes identified in these BlastFrost queries contain a single nucleotide variant in *gyrA* (89.7%) (Fig. 5D). Variants in *gyrB* (1.8%) and *parE* (0.26%) were also found but they were less common, and were normally present together with *gyrA* mutations in the same genomes (Fig. 5C).

**Figure 5:**
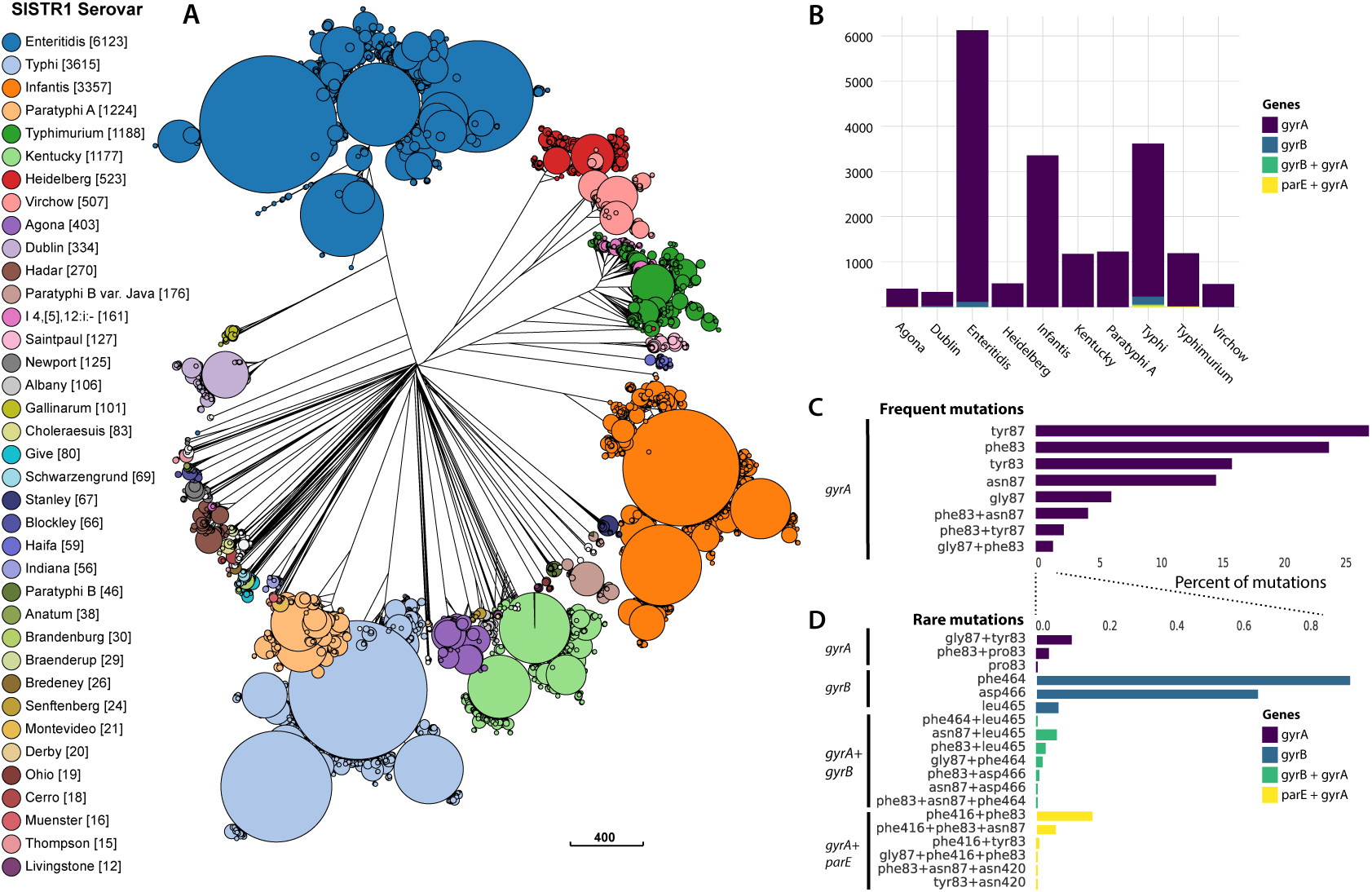
20, 490 *Salmonella* genomes containing QRDR variants of genes *gyrA, gyrB* and *parE* that are associated with fluoroquinolone resistance [26]. (A) Neighbor-Joining tree based on cgMLST distances (visualized in GrapeTree [31]), colored according to the *Salmonella* serovar in EnteroBase [30] according to SISTR1 [28]. (B) Distribution of mutated genes by serovar for the ten most frequent serovars in part A. (C+D) Percentage of individual nucleotide mutations, and their combinations by frequent (C) and rare (D) mutations.

EnteroBase also contains numerous genomes which do not contain these QRDR mutations. The relative proportions of genomes with and without those QRDR mutations from the most frequent serovars are illustrated in Figure 6. Serovars Paratyphi A or Typhi show the largest proportion of strains with resistance mutations. Interestingly, almost all fluoriquinolone-resistants strains of serovar Kentucky belong to only one of the multiple genetic clusters that are associated with this polyphyletic serovar [1, 2].

**Figure 6:**
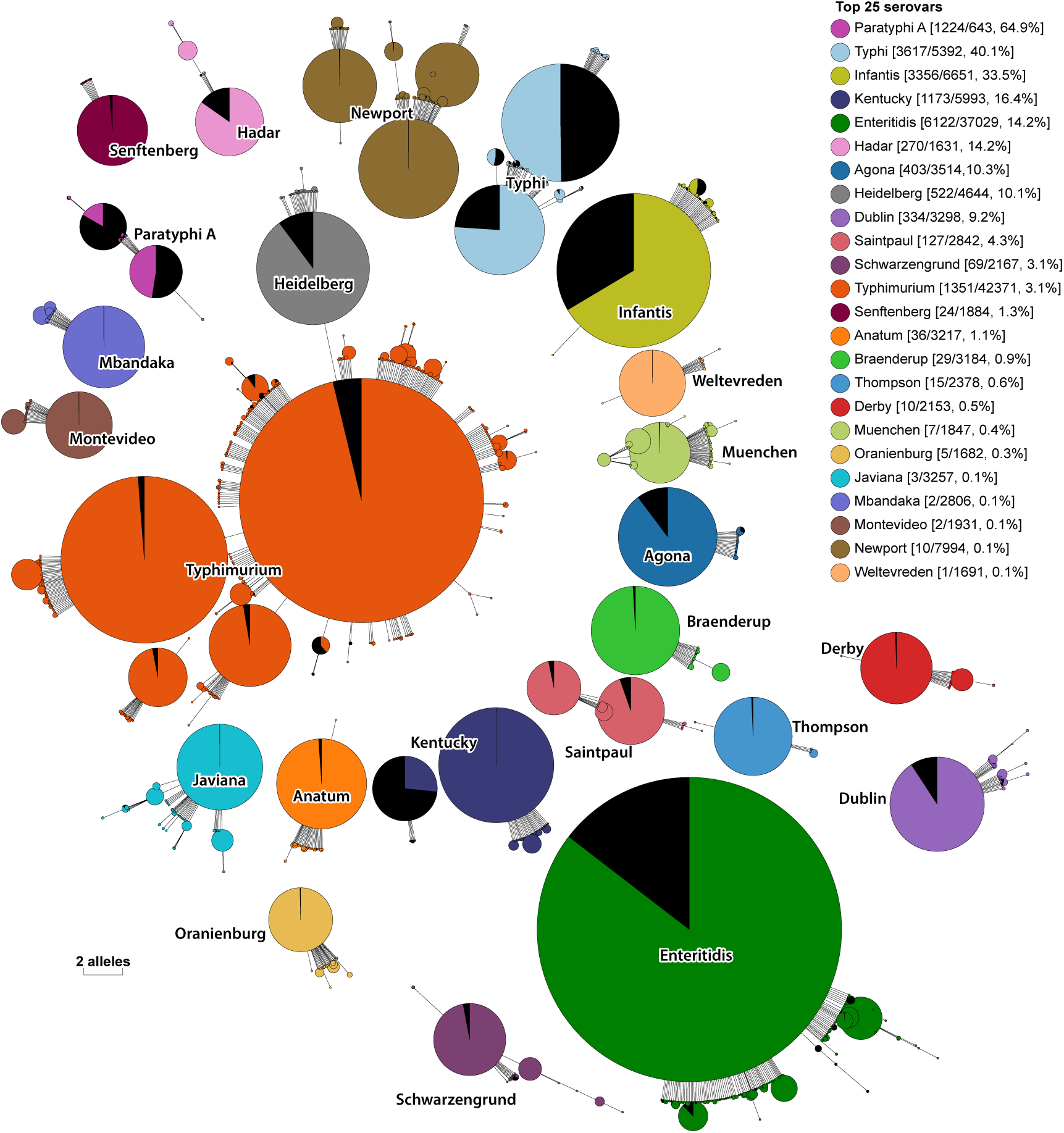
Proportion of *Salmonella* genomes containing QRDR mutation (black) among all genomes in top 25 HC900 clusters (colored) in EnteroBase [30]. EnteroBase uses hierarchical single-linkage clustering based on cgMLST distances to derive clusters at different cut-off levels. A cut-off of 900 has been identified to correlate well with serovar predictions, hence clusters are labelled by their majority serovar (EnteroBase conversion table: https://enterobase.readthedocs.io/en/latest/HierCC_lookup.html). The Neighbour-Joining tree of all genomes in these clusters is based on 7-gene MLST distances, and visualized in GrapeTree [31]. For each cluster, the legend indicates the number of genomes with and without the QRDR mutation, and its percentage.

These analyses took 25 minutes to load the Bifrost graph into memory, and 3.5 hours to extract all subgraphs using 8 threads. BlastFrost used a maximum of 160GB RAM for these analyses.

## 3 Discussion and Conclusions

BlastFrost implements a highly efficient algorithm for querying de Bruijn graphs. It complements the very computationally efficient Bifrost, which calculates compacted and colored graphs that support scaling such analyses to 100,000s of closely related bacterial genomes. Practical applications of these two methods are also greatly facilitated by the existence of structured sequence databases of closely related bacteria such as EnteroBase. Furthermore, visualisation of the genetic relationships among the query hits is also facilitated by the genotyping based on legacy or core genome MLST, which is also provided by EnteroBase [30]. The combination of Bifrost, BlastFrost and EnteroBase has the potential to rapidly reveal numerous features of genomic diversity that were previously not readily possible.

MLST schemes, even whole genome MLST, are inherently limited, because they are based on a fixed selection of genes that were present in an initial, representative set of genomes. However, many bacterial genera are associated with open pan-genomes [21], whose content continues to increase with each additional genome that is sequenced [29]. These novel sequences will not be included in MLST schemes. Therefore, it is important to emphasise that BlastFrost and Bifrost are not dependent on MLST or on genomic annotations, but can handle any collection of closely related genomic assemblies. BlastFrost can summarise diversity within large regions such as genomic islands. It can identify variants of any sequence of interest, which can be rapidly analysed to find single nucleotide polymorphisms.

We compared the speed and memory requirements of BlastFrost and Bifrost for large genomic data sets with the state of the art tools BIGSI and minimap2. BIGSI can handle genomic analyses between diverse bacterial genomes, whereas BlastFrost is less suitable for indexing and querying diverse sequence collections such as RefSeq or SRA. However, for closely related genomes, such as those within a single bacterial genus, BlastFrost is considerably faster than BIGSI, and requires less memory for up to 1400 sequence queries. Similarly, BlastFrost is much faster than minimap2 for closely related genomes, and also requires less memory. These computational efficiencies did not sacrifice accuracy. BlastFrost has good precision and sensitivity for sequences that are at least 90% identical and over 400bp in length.

The identification of genomic islands or individual nucleotides associated with antimicrobial resistance genes is enabled by BlastFrost because of the explicit graph data structure in Bifrost which supports graph traversal and extraction of sequences that extend beyond the k-mers that were used for querying. As a result, given a Bifrost graph, genomic islands or nucleotide variants can be identified among 100,000s of genomes in a matter of minutes. The Bifrost API freely supports annotation of nodes in the graph, including annotating unitigs with additional data. In future extensions, BlastFrost should be able to extract local synteny from graphs whose unitigs are annotated with genome coordinates and/or gene annotations. Such information could also be used to reconstruct genomic rearrangements.

BlastFrost is not a general replacement for calling SNPs because its precision suffers with increasing genetic diversity and reduced sequence length. However, it might have the potential for incorporation into approaches to detection of antimicrobial resistance in combinations of databases of AMR genes such as CARD [14] and AMRfinder [10] and genomic sequence collections such as EnteroBase.

## Acknowledgements

This project was supported by the Wellcome Trust (202792/Z/16/Z). GH was supported by the Icelandic Research Fund Project grant number 152399-053. We thank Zhemin Zhou, Jane Charlesworth and Páll Melsted for their helpful feedback during the development of BlastFrost.

