## Supplemental Material for "BlastFrost: Fast querying of 100,000s of bacterial genomes in Bifrost graphs"

### A Data

All genome assemblies used in the analyses in this paper were downloaded from EnteroBase. For each dataset, a file containing all EnteroBase barcodes for these assemblies has been stored under `https://github.com/nluhmann/BlastFrost/data`. The wgMLST scheme for *Salmonella* in EnteroBase can be downloaded from `https://enterobase.warwick.ac.uk/species/senterica/download_data`. The sequences for SPI-2 genes were downloaded from the Virulence Factor Database at `http://www.mgc.ac.cn/cgi-bin/VFs/genus.cgi?Genus=Salmonella`. The accession numbers for genes *gyrA*, *gyrB* and *parE* are

### B Methods

#### B.1 Data structures

We consider as input a Bifrost graph of a set of genomes $G$, and a set of query sequences $Q$. As EnteroBase contains a standardised assembly pipeline for all short read dataset added to the database (either through user upload or fetched from public databases), we assume all genomes in $G$ to be draft assemblies consisting of a set of contigs. However, the following is equally valid for graphs build on sets of sequencing reads.

**colored DBG (cDBG)** Define $K_g$ as the set of unique k-mers (words of length k) over an alphabet $\Sigma$ present in a genome $g \in G$. We denote $K = \cup_{g \in G} K_g$. Let $C$ be a set of colors and $c_g \in C$ be a unique color assigned to a genome or a cluster of genomes respectively. Define $dBG(V, E)$ as a de Bruijn graph, with the set of vertices $V$ representing all k-mers in $K$, i.e. $V = \cup_{g \in G} K_g$. The edges in the graph are given implicitly by two k-mer sequences overlapping in exactly in $k - 1$ positions, i.e. $k_1[2 : n] = k_2[1 : n - 1]$. In a colored de Bruijn graph $cdBG(V, E)$, each vertex $v \in V$ is labeled with a set of colors $c_v \subset C$, consisting of all $c_g$ for which $k_v \in g$.

The complexity of the colored dBG can be reduced by compacting maximal non-branching paths into single vertices, so called unitigs. We denote the compacted version of the colored de Bruijn graph as $ccDBG$. Note that while each compacted vertex represents a maximal non-branching path in terms of sequence, depending on sequencing coverage or assembly quality, we do not assume all $k$-mers compacted into a single vertex to be labeled with the same set of colors.

**Bifrost graphs** We rely on a recently developed data structure for compacted and colored dBGs named Bifrost [12]. Bifrost is a parallel and memory efficient tool enabling the direct construction of the compacted de Bruijn graph without producing the intermediate uncompacted de Bruijn graph. Its software library features a broad range of functions such as graph traversal, querying and editing while automatically preserving the compaction property. Bifrost makes full use of a dynamic index to update the graph with additional genomes and efficiently colour each $k$-mer of the graph with the set of genomes in which it occurs.

As our following method relies on functions defined in the Bifrost API, we will refer to the graph required as input as Bifrost graphs. All Bifrost graphs underlying the analysis and evaluation in Section 2 have been build with Bifrost parameters $k = 31$ and $m = 21$, and options $-r$ for Bifrost reference mode and $-c$ to include the coloring of the graph.

We built Bifrost graphs for all draft assemblies in EnteroBase up to July 2019. Table 1 shows the time and memory requirements for the initial built of Bifrost graphs for the two largest databases in EnteroBase September 2018. Since then, the *Salmonella* Bifrost graph has been updated to now contain $190,209$ strains. The graph and colors file for this graph have a total uncompressed size of 158.5GB on disk.

Table 1: Time and memory requirements to build Bifrost graphs for draft assemblies of *Salmonella* and *Escherichia* in Enterobase as of September 2018.

| | | *Salmonella* | *Escherichia* |
|---|---|---|---|
| Input | # assemblies | 160k | 72k |
| | size | 1.5 TB | 728 GB |
| Bifrost | time | 4 days, 15h | 1 day, 15h |
| | memory | 147 GB | 102 GB |
| Output | file size | 125 GB | 86 GB |
| | unitigs | >30M | >32M |
| | edges | 86M | 93M |

For the collection of 736 *Yersinia* draft assemblies, we compared the build time and memory requirements for BIGSI and Bifrost on standard parameters (see Figure 7). As *Yersinia* is a rather conserved bacterial genus, Bifrost is able to index these genomes more efficiently than BIGSI.

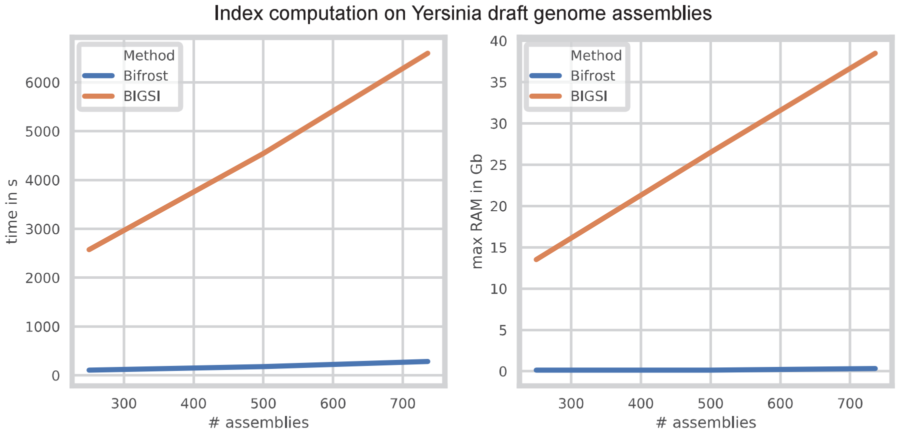


Figure 7: Benchmark comparison of Bifrost and BIGSI. We built both the Bifrost graph and BIGSI index on subsets of 736 draft assemblies of *Yersinia* representative strains.

#### B.2 K-mer query in BlastFrost

BlastFrost accepts as input a colored Bifrost graph, consisting of a GFA file describing the graph (parameter $-g$) and a binary file representing the color information for the graph (parameter $-f$). Query sequences are specified with parameter $-q$ in multiple FASTA format. As optional parameters, the number of threads (default 1) and distance parameter $d$ (default 1) can be specified. The default analysis will output the presence/absence analysis for each query, while parameter $-e$ enables subgraph extraction described below.

Given a set of query sequences $Q$, we apply the same value $k$ used to build the underlying Bifrost graph to derive a set of unique k-mers $K_q$ for each $q \in Q$. As the search for each query against the graph is considered independently, we simplify the following description by describing the search for a single query $q$.

We load the pre-computed Bifrost graph back into memory using the Bifrost API function respectively.

For each $k \in K_q$ ordered in ascending position of their occurrence in $q$, we query the Bifrost graph for the unitig the $k$-mer appears in. This results in a binary hit sequence $b_c$ for each color hit by any $k$-mer in the query (1 indicating the $k$-mer has been found in a unitig colored by $c$, 0 indicating absence). Note that a $k-mer$ compacted in a unitig does not have to contain all colors present in the whole unitig due to e.g. sequencing coverage or assembly quality. In order to reduce computation steps, we assume two query $k$-mers located on the same unitig to be labeled with the same set of colors.

In order to increase the sensitivity of the k-mer based query search, we implement an inexact search by additionally querying for matches of $k$-mers in the $d$-neighborhood of a query k-mer $k$, with $d$ being an input distance parameter that can be chosen by the user:

$N_d^{\delta}(k) = \{k_2 : \delta(k, k_2) \leq d \wedge |k_2| = k\}$

Note that the considered neighborhood is restricted to sequences of length $k$ as this value is fixed to query the Bifrost graph. Depending on the size of the query and the value for $d$, this will increase the number of queried $k$-mers significantly. Our experiments however showed only small increase in run time for queries up to 2000bp and $d = 2$, while the sensitivity increased.

Given a hit sequence $b_c$, we estimate an alignment score for the query against color $c$ by measuring the length of 0 runs. We assume that a single substitution will cause $k$ $k$-mers to be missed, hence we divide the number of 0s in contiguous runs by $k$. We use a standard score for matches of 1 and mismatches of $-2$. From this estimated alignment score, we calculate a p-value using the BLAST formula.

**Algorithm 1 BlastFrost**: k-mer presence/absence query

**Require:** BlastFrost graph $G$, query sequence $q$, distance $d$
- $K_q \leftarrow k - mers(q)$
- **for** $i \in [1..|K_q|]$ **do**
  - $u \leftarrow G.find(K_q[i])$
  - **if** $u.empty() d > 1$ **then**
    - $N_d^\delta(k) = \{k_2 : \delta(k, k_2) \leq d \wedge |k_2| = k\}$
    - **for** $k'$ $inN_d^\delta$ **do**
      - $u \leftarrow G.find(K_q[i])$
    - **end for**
  - **end if**
  - $C_u \leftarrow u.getUnitigColors()$
  - **for** $c \in C_u$ **do**
    - $b_c[i] = 1$
  - **end for**
- **end for**

**Subgraph extraction** For each query $q \in Q$, the previous step identified a list of unitigs that were found in the $k$-mer search for a subset of colors $C_q \in C$. We define $U_c$ as the set of unitigs that have been found for color $c$. For each $c \in C_q$, we aim to complete a path in the graph that is pre-sketched by the k-mer hits. A perfect hit of a query $q$ against a color $c$ in the graph will result in an already complete path, whereas variation between the query and the graph genome will cause gaps in the graph path respectively.

In order to avoid unnecessary graph traversal, we first cluster colors in $C_q$ with the same unitig sets, assuming that multiple colors can represent the same query variant and therefore follow the same paths in the graph. However, colors being grouped together initially can diverge during any step of the graph traversal and will require a subsequent separate path traversal.

For each cluster of colors, we start with the first listed unitig $u$. Note that this unitig does not have to contain first k-mer in the query. We follow successors of $u$ in the graph in a depth-first search, adding them to the unitig list and comparing against the current cluster of colors, until we traverse a unitig that is already in the set of identified unitigs for this cluster of colors or until we hit a extension limit. If this limit is hit, we remove all colors from the list of query hits and do not report a subgraph. The extension ends successfully if we find the last unitig identified by a k-mer hit.

Concatenating the extracted path unitigs, considering their overlap of length $k-1$ at their ends, gives the sequence underlying the identified path in the graph. The length of this sequence can be shorter than the initial query,as the first or last unitig do not have to align with the first or last k-mer in the query. We subsequently extend the path to the left or right until the length of the path sequence is equal to the query length or until the path for the current color cluster becomes ambiguous, i.e. there are more than one preceding or succeding unitigs found for the current color.
